# CDK2 activity determines the timing of cell-cycle commitment and sequential DNA replication

**DOI:** 10.1101/2020.06.24.169409

**Authors:** Sungsoo Kim, Alessandra Leong, Chellam Nayar, Minah Kim, Hee Won Yang

## Abstract

To enter the cell cycle, mammalian cells must cross a point of no return (the commitment point), after which they proceed through the cell cycle regardless of changes in external signaling. This process is tightly regulated by the cyclin-dependent kinases (CDKs) and downstream molecules such as retinoblastoma (Rb). Here we show that CDK2 activity coordinates the timing of cell-cycle commitment and DNA replication. CDK4/6 activation initiates Rb phosphorylation and E2F activity, causing a gradual increase in CDK2 activity. Once CDK2 activity reaches a threshold level, CDK2 triggers the commitment point by maintaining Rb phosphorylation and subsequently initiates DNA replication. While the timing of the commitment point is tightly coupled with DNA replication, our experiments, which acutely increased CDK2 activity, suggest that the timing of the commitment point is before DNA replication. These findings highlight how cells utilize a safety mechanism to maintain genome stability by protecting against incomplete DNA replication.

## Introduction

Entry into the cell cycle is tightly regulated in multicellular organisms, and dysregulation of this process can lead to cancer^1^ and degenerative diseases^2^. Extracellular signaling controls cell-cycle entry until cells irreversibly commit to the cell cycle (the commitment point) to ensure faithful DNA replication^3,4^. In mitogen-starved cells, Pardee first defined a specific point in late G1, the restriction point, where cells commit to the cell cycle regardless of mitogen availability^5^. In addition to mitogenic signaling, stress signals including DNA damage and osmotic stress can also regulate cell-cycle entry^6–9^. In asynchronously cycling cells, recent studies demonstrated that while cells can commit to the next cell cycle in G2 phase of mother cells with respect to mitogen availability^10^, stress can reverse cell-cycle entry until inactivation of the anaphase-promoting complex/cyclosome-Cdh1 (APC/C^Cdh1^) at the G1/S transition^8,11^.

Mechanistically, the retinoblastoma (Rb)/E2F pathway is central to the regulation of the commitment point. Mitogens upregulate the expression of cyclin D, resulting in activation of CDK4 and CDK6 (CDK4/6). CDKs activity regulates the ability of Rb to bind E2F by phosphorylating 15 putative CDK phosphorylation sites on Rb^12^. During quiescence, Rb serves as a transcriptional co-repressor that binds E2F proteins and represses their transcriptional activities^4,13^. It has been proposed that partial phosphorylation of Rb by CDK4/6 is sufficient to disrupt the Rb/E2F interaction and initiate the cell cycle^4,13–18^. However, a recent report showed that CDK4/6 mono-phosphorylates Rb, which can exist in un-, mono-, and hyper-phosphorylated forms with no evidence of partial phosphorylation^19^. Considering that phosphorylation of individual residues causes major changes to the Rb structure^20–22^, another recent study revealed that distinct mono-phosphorylated forms of Rb alter the composition of the endogenous Rb complexes and induce different transcriptional outputs^23^. A subsequent study demonstrated, however, that CDK4/6 activity alone is sufficient and required for full phosphorylation of Rb (hyper-phosphorylation) throughout the G1 phase^24^, indicating the complexity of CDK4/6 regulation of Rb.

Once Rb is inactivated, E2F transcription factors induce cyclin E to activate CDK2. Previous studies showed that CDK2 activation induces Rb hyper-phosphorylation and generates a CDK2-Rb positive feedback loop, triggering irreversible Rb phosphorylation^4,13–15,25^. A recent study further characterized that CDK2-Rb feedback is engaged at the onset of S phase in both asynchronously cycling and mitogen-starved cells^24^. In addition, the E3 ubiquitin ligase APC/C^Cdh1^ controls cell-cycle progression by promoting the degradation of key cell-cycle regulators, such as cyclin A and geminin^26,27^. E2F transcription factors also induce expression of Emi1 which inactivates APC/C^Cdh1(11,28,29)^. Another study revealed that irreversible APC/C^Cdh1^ inactivation at the G1/S transition controls the commitment point with respect to stress^8^. Double-negative feedback between APC/C^Cdh1^ and Emi1 underlies the molecular mechanism of irreversible inactivation of APC/C^Cdh1(11)^. These studies suggest that cells utilize different feedback mechanisms to trigger the commitment point depending on mitogenic and stress signaling. Since both APC/C^Cdh1^-Emi1 and CDK2-Rb feedbacks engage at the G1/S transition^8,24^, inactivation of CDK4/6 before the initiation of S phase can reverse cell-cycle entry. Indeed, it was demonstrated that while CDK4/6 activity can be sustained after mitogen withdrawal^24^, the induction of various stress signals can rapidly suppress CDK4/6 activity^9^, possibly explaining different temporal cell-cycle commitment points with respect to mitogens or stress. However, it is unknown how distinct APC/C^Cdh1^-Emi1 and CDK2-Rb feedbacks coordinate cell-cycle commitment and the transition into S phase. Additionally, given that high CDK2 activity can phosphorylate Rb at the onset of S phase^9^, it is unclear whether other mechanisms in S phase are required to initiate CDK2-Rb feedback.

Here, we investigated regulation of the Rb/E2F pathway by CDK4/6 and CDK2 in relation to the commitment point by using a live-cell reporter for CDK4/6 activity^9^. We propose that CDK2-Rb feedback is the primary signaling network regulating cell-cycle commitment with respect to mitogenic and DNA damage signaling. Our results show that CDK4/6 activation initiates Rb phosphorylation and subsequently E2F activity. Then, a threshold level of CDK2 activity coordinates the timing of cell-cycle commitment and DNA replication, controlling the transition into S phase. APC/C^Cdh1^ inactivation also potentiates cell-cycle commitment by upregulating CDK2 activity. We found that while this cell-cycle commitment is typically triggered at the onset of S phase, an increase in CDK2 activity is sufficient to initiate CDK2-Rb feedback before DNA replication. Together, our studies suggest that Rb phosphorylation regulated by a CDK2 activity threshold is the mechanism of irreversible cell-cycle entry.

## Results

### Rb phosphorylation by CDK4/6 can trigger the initiation of E2F activity

To explore how CDK4/6 and CDK2 coordinate the Rb/E2F pathway in relation to APC^Cdh1^ inactivation, we used live-cell reporters for CDK4/6, CDK2, and APC/C ^Cdh1^ activities^8–10,30,31^ in human epithelial MCF-10A cells (**Fig. 1a**). In the same cells, we performed immunostaining to measure Rb phosphorylation at S807/811 and mRNA fluorescence *in situ* hybridization (FISH) to measure the E2F target gene, E2F1. We note that the CDK2 reporter generally measures the collective activity of cyclin E/A-CDK1/2 complexes^32^. The CDK4/6 reporter contains a C-terminal fragment of Rb protein (a.a. 886 − 928) and measures Rb phosphorylation by CDK4/6 and CDK2 (**Fig. 1b**). To account for CDK2 activity in S and G2 phase that can be captured by the CDK4/6 reporter, we previously derived a corrected CDK4/6 activity by subtracting a calculated fraction (35%) of the CDK2 reporter signal from the CDK4/6 reporter signal in MCF-10A cells^9^.

**Figure 1.**
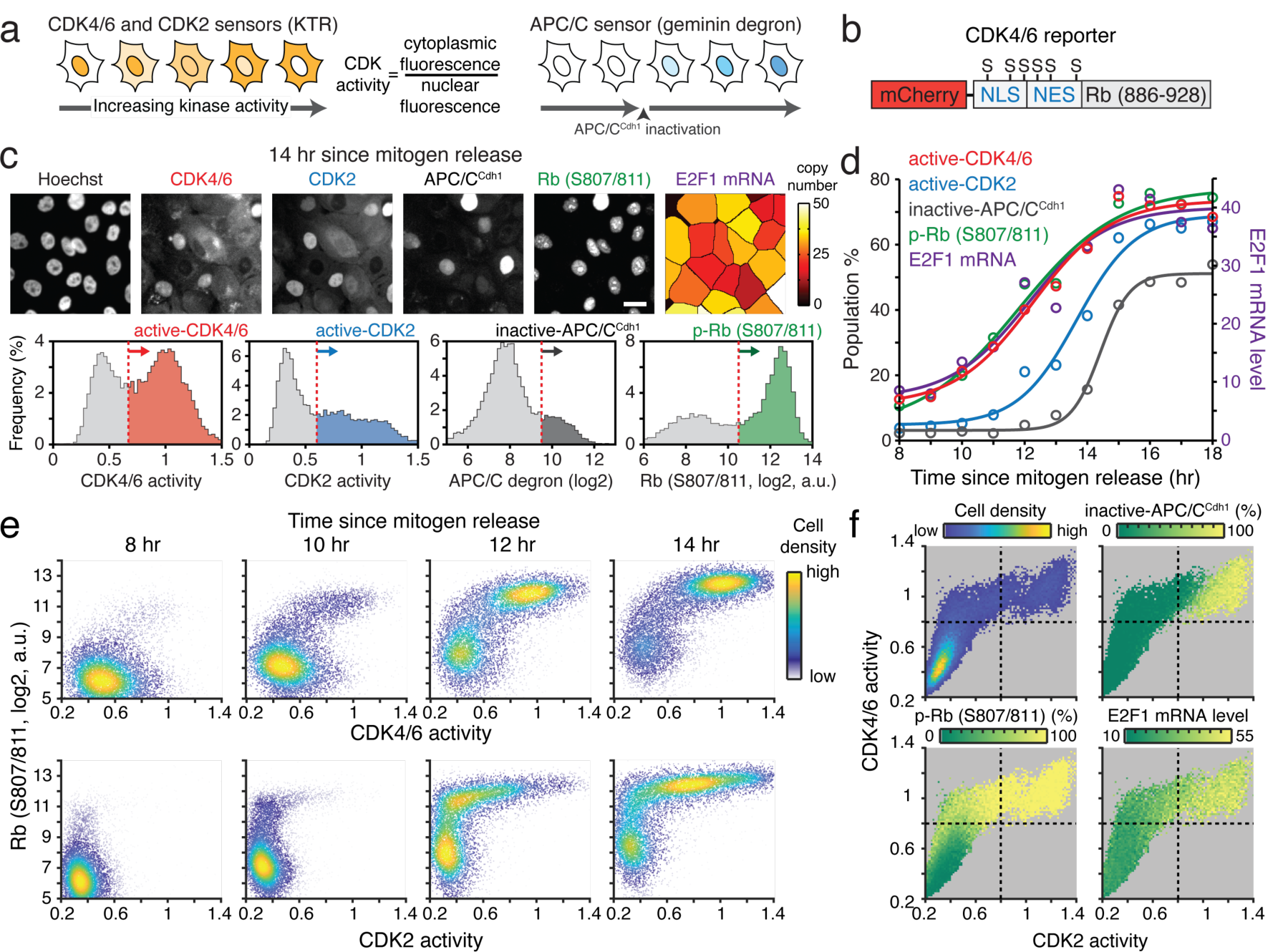
Activation of CDK4/6 is tightly coupled with Rb phosphorylation and E2F1 mRNA induction. **(a)** Schematics of CDK4/6, CDK2 reporters (left) and APC/C^Cdh1^ reporter (right). Active CDK4/6 and CDK2 phosphorylate and induce translocation of reporters to the cytoplasm. As APC/C^Cdh1^ is inactivated, intensity of the reporter starts increasing. **(b)** Schematic of the CDK4/6 reporter. **(c)** Top, representative images of CDK4/6, CDK2, and APC/C^Cdh1^ reporters, Rb (S807/811) immunostaining, and mRNA FISH for E2F1 mRNA in the same MCF-10A cells. Scale bar is 20 µm. Bottom, histograms of CDK4/6, CDK2, and APC/C^Cdh1^ activities and phosphorylated Rb (p-Rb) at S807/811. Red dotted line indicates a threshold used to classify active CDK4/6 and CDK2, inactive APC/C^Cdh1^, and p-Rb populations. Cells were mitogen-released for 14 hr prior to fixation. **(d)** Scatter plot of the percentage of cells with active CDK4/6, active CDK2, inactive APC/C^Cdh1^, and p-Rb at S807/811 in MCF-10A cells as a function of time after mitogen release. E2F1 mRNA level reflects averaged value over time. Solid line is sigmoidal best-fit line. **(e)** Single-cell correlation of p-Rb at S807/811 versus CDK 4/6 activity (top) and CDK2 activity (bottom) in MCF-10A cells at various time points after mitogen release (8, 10, 12 and 14 hr). Cell density is color-coded. **(f)** Three-dimensional activity map of CDK4/6 versus CDK2 activity in MCF-10A cells where cell density, the percentage of inactivated APC/C^Cdh1^, the percentage of p-Rb at S807/811, and averaged E2F1 mRNA level are color-coded. Black dotted lines represent CDK4/6 activity onset and CDK2 activity onset.

In asynchronously cycling cells after mitosis, cells continuously trigger CDK4/6 activation, Rb phosphorylation, and high E2F activity and immediately start CDK2 activation to continue cell-cycle entry^9,33^. Therefore, to explore the initiation of the Rb/E2F pathway by CDK4/6 and CDK2, we synchronized cells in quiescence (G0) by mitogen starvation for 48 hours. Then, we stimulated cells with mitogens (hereafter mitogen release) and fixed them at various time points. Based on the distributions of the CDK activities, intensity of the APC/C^Cdh1^ reporter, and Rb phosphorylation levels, we classified populations for active CDK4/6 and CDK2, inactive APC/C^Cdh1^, and Rb phosphorylation at S807/811 (**Fig. 1c**). When we plotted the percentages of cells in each population and analyzed the averaged E2F1 mRNA level as a function of time after mitogen release, we found that CDK4/6 activation, Rb phosphorylation at S807/811, and E2F transcription activity were tightly coordinated before CDK2 activation and APC/C^Cdh1^ inactivation (**Fig. 1d**). Furthermore, at the single-cell level, CDK4/6 activation and Rb phosphorylation at S807/811 were strongly correlated, while CDK2 activity started increasing after both Rb phosphorylation at S807/811 and onset of CDK4/6 activity (**Fig. 1e**). To classify cells based on CDK4/6 and CDK2 activities, we analyzed three-dimensional activity maps of CDK4/6 versus CDK2 where the percentages of inactive APC/C^Cdh1^, Rb phosphorylation at S807/811, and E2F1 mRNA levels are color-coded. These plots showed that activation of CDK4/6 was tightly correlated with Rb phosphorylation at S807/811 and increased E2F1 mRNA levels (**Fig. 1f**). APC/C^Cdh1^ was inactivated only in the cells with both high CDK4/6 and CDK2 activities (**Fig. 1f**). Together, our data indicate that Rb phosphorylation at S807/811 by CDK4/6 is tightly correlated with E2F activity.

We next performed multiplex staining^34^ to sequentially measure multiple sites of Rb phosphorylation at S807/811, T373, S608, and S780 in the same MCF-10A cells (**Fig. 2a**). We confirmed that all Rb-phosphorylation sites measured by four different antibodies were closely correlated at the single-cell level^24^ (**Supplementary Fig. 1a**). By analyzing three-dimensional activity maps of CDK4/6 versus CDK2 color-coded for the percentage of Rb phosphorylation, we found that CDK4/6 activity was tightly coupled with all four measured Rb phosphorylation sites at S807/811, T373, S608, and S780 (**Fig. 2b**). Furthermore, E2F1 mRNA levels started increasing at the boundary of Rb phosphorylation at S807/811 (**Fig. 2c**), suggesting that Rb phosphorylation by CDK4/6 may initiate E2F activity. In a similar manner, we observed that CDK4/6 activation was tightly coupled with Rb phosphorylation at S807/811 and E2F activation in asynchronously cycling and passage-limited human lung fibroblasts (HLF) and human foreskin fibroblasts (HS68), as well as hTERT immortalized human foreskin fibroblasts (BJ-5ta) (**Fig. 2d, e** and **Supplementary Fig. 1b**). These results indicate that regulation of Rb phosphorylation at S807/811 and E2F activity is conserved across several primary and immortalized cell types.

**Figure 2.**
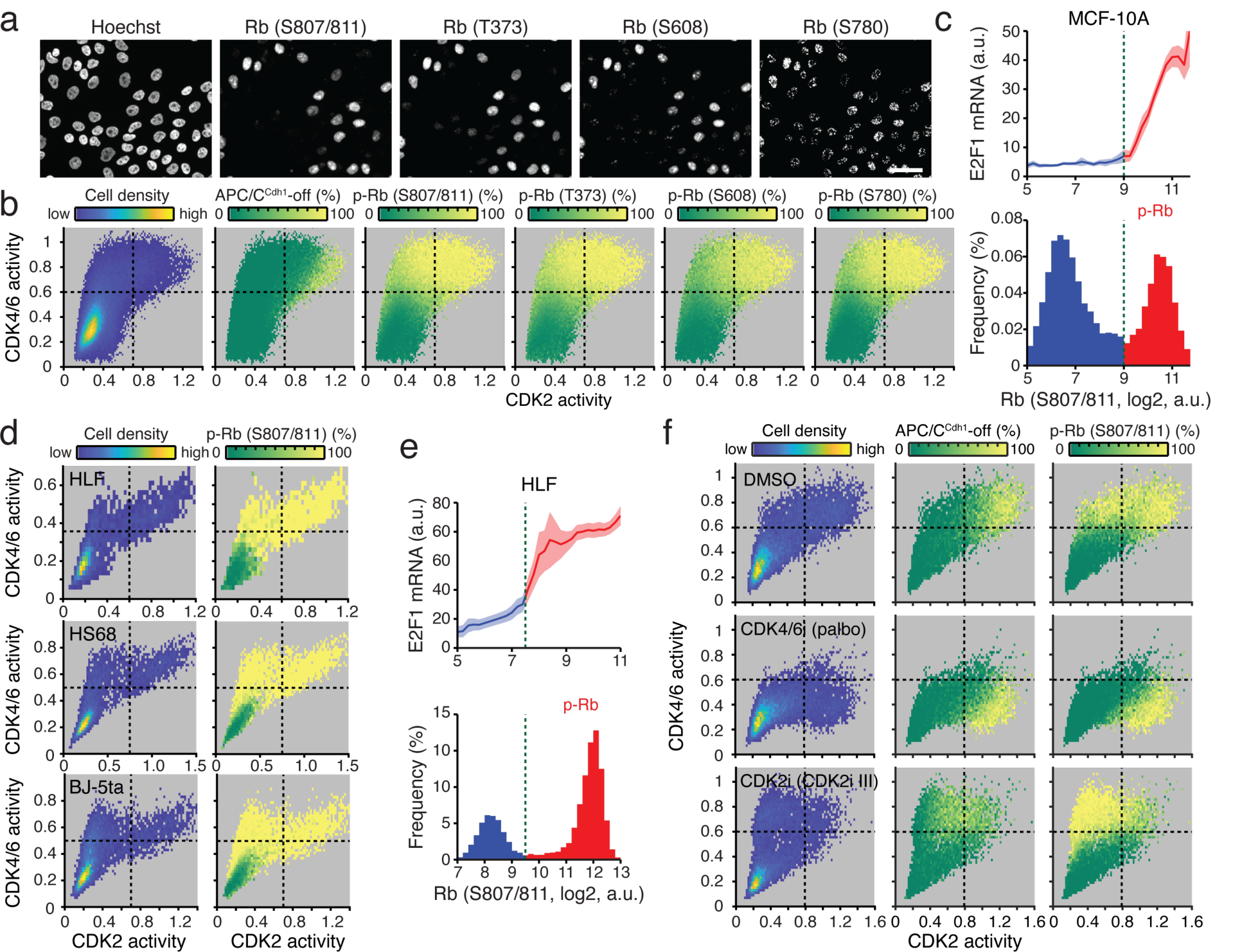
CDK4/6 activation triggers Rb phosphorylation at multiple sites. **(a)** Representative images of multiplexed staining of Rb (S807/811, T373, S608, S780) in MCF-10A cells. Scale bar is 20 µm. **(b)** Three-dimensional activity map of CDK4/6 versus CDK2 activity where cell density, the percentage of inactivated APC/C^Cdh1^, and the percentage of p-Rb at sites S807/811, T373, S608, and S780 are color-coded. MCF-10A cells were mitogen-released for 12 hr prior to fixation. Black dotted lines represent CDK4/6 activity onset and CDK2 activity onset. **(c)** Top, E2F1 mRNA levels across the distribution of p-Rb at S807/811 in MCF-10A cells. Data are mean ± 95% confidence intervals (*n* = 11,741 cells). Bottom, histogram of p-Rb at S807/811. **(d)** Three-dimensional activity map of CDK4/6 versus CDK2 activity in HLF, HS68, and BJ-5ta cells (top to bottom). Cell density and the percentage of p-Rb at S807/811 are color-coded. Black dotted lines represent CDK4/6 activity onset and CDK2 activity onset. **(e)** Top, E2F1 mRNA levels across the distribution of p-Rb at S807/811 in HLF cells. Data are mean ± 95% confidence intervals (*n* = 18,861 cells). Bottom, histogram of p-Rb at S807/811. **(f)** Three-dimensional activity maps of CDK4/6 versus CDK2 activity under the control condition, CDK4/6 inhibition, and CDK2 inhibition (top to bottom). The percentage of p-Rb at S807/811, cell density and APC/C^Cdh1^ are color-coded. After mitogen release for 12 hr, MCF-10A cells were treated with Palbociclib (1 µM) or CDK2i III (60 µM) for 1 hr prior to fixation. Black dotted lines represent CDK4/6 activity onset and CDK2 activity onset.

To further investigate the effect of CDK4/6 on Rb phosphorylation, we used CDK4/6 inhibitors (palbociclib, abemaciclib, and ribociclib). In light of a recent paper showing that long-term incubation of CDK4/6 inhibitors (48 hours) can suppress CDK2 activity by redistributing the CDK inhibitor, p21^(35)^, we acutely treated cells with CDK4/6 inhibitors for 1 hour. All CDK4/6 inhibitors reversed Rb-phosphorylation at S807/811, T373, S608, and S780 in cells with high CDK4/6 and low CDK2 activities (**Fig. 2f** and **Supplementary Fig. 2**). Consistent with a previous report showing high CDK2 activity is required to maintain Rb phosphorylation^24^, we found that Rb phosphorylation was resistant to CDK4/6 inhibition in cells with high CDK2 activity and inactive APC/C^Cdh1^. In addition, high CDK4/6 activity was enough to maintain the Rb phosphorylation sites after acute treatment with CDK2 inhibitors (roscovitine and CDK2i III (CVT-313)). Together, our data are consistent with previous reports showing that CDK4/6 either randomly mono-phosphorylates or hyper-phosphorylates Rb^19,24^. In addition, our finding suggests that Rb phosphorylation mediated by CDK4/6 activity initiates E2F activity.

### Rb phosphorylation by CDK4/6 is sufficient to initiate E2F activity

To explore the kinetics of Rb phosphorylation and E2F activity regarding CDK4/6 and CDK2 activation, we performed live-cell imaging to monitor CDK4/6, CDK2, and APC/C^Cdh1^ activities and fixed-cell analysis to measure Rb phosphorylation and mRNA levels of E2F target genes in MCF-10A cells. After mitogen release, cells either activated CDK4/6 to initiate proliferation (CDK4/6^high^) or maintained low CDK4/6 activity and subsequently remained in quiescence^9^ (CDK4/6^low^) (**Supplementary Fig. 3a**). CDK4/6^high^ cells consequently induced CDK2 activation and APC/C^Cdh1^ inactivation and exhibited substantial heterogeneity in the timing of CDK4/6 activation (**Fig. 3a** and **Supplementary Fig. 3a**), allowing us to measure Rb phosphorylation and E2F target genes at various time points after CDK4/6 activation. By mapping fixed-cell data back to live-cell data at the single-cell level, we compared Rb phosphorylation levels between CDK4/6^high^ and CDK4/6^low^ cells and found that all five Rb phosphorylation sites were highly correlated with CDK4/6^high^ cells (**Supplementary Fig. 3b**).

**Figure 3.**
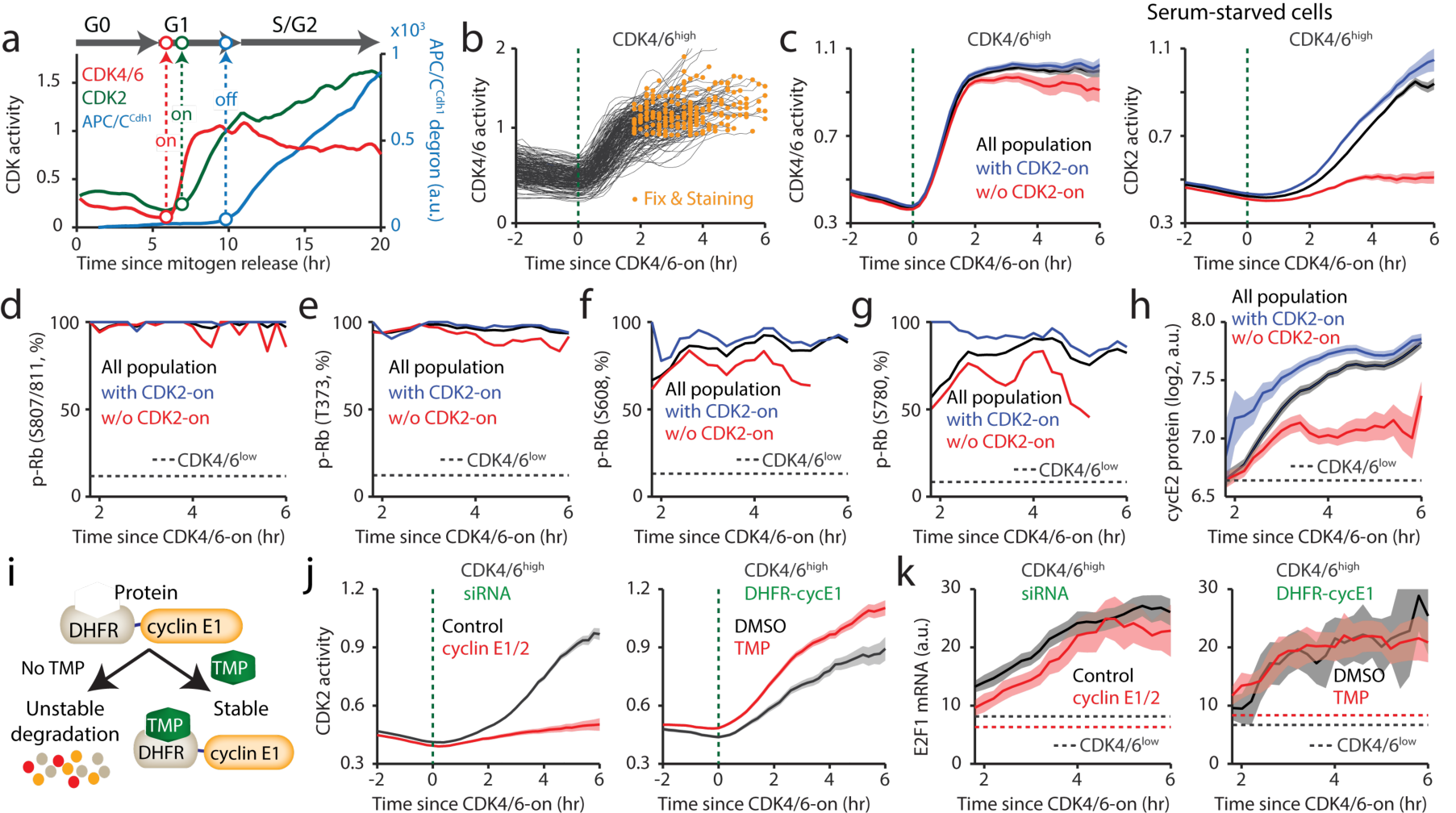
CDK4/6 activation triggers immediate Rb phosphorylation and E2F activation. **(a)** Representative time course analysis of CDK4/6 activity, CDK2 activity, and APC/C^Cdh1^ inactivation labeled with cell-cycle phase and demarcation of CDK activation and APC/C^Cdh1^ inactivation point. **(b)** Single-cell traces of CDK4/6 activity aligned to the time of CDK4/6 activation in CDK4/6^high^ cells. To create more heterogeneity, MCF-10A cells were mitogen-released for various durations (9, 10, and 11 hr) prior to fixation. Orange dots correspond to fixation point. Green dotted line indicates CDK4/6 activity onset. **(c)** Averaged CDK4/6 activity traces (left) and CDK2 activity traces (right) aligned to CDK4/6 activation in MCF-10A cells. Black, blue, and red lines correspond to all cells, cells with CDK2 activity, and cells without CDK2 activity, respectively. Green dotted line indicates CDK4/6 activity onset. Data are mean ± 95% confidence interval (All population, *n* = 1,917 cells; with CDK2 activity, *n* = 879 cells; w/o CDK2 activity, *n* = 1,038 cells). **(d−g)** Percentage of p-Rb at sites S807/811 (d), T373 (e), S608 (f), and S780 (g), as a function of CDK4/6 activation in MCF-10A cells. Dotted line indicates percentage of p-Rb in CDK4/6^low^ cells. Black, blue, and red lines correspond to all cells, cells with CDK2 activity, and cells without CDK2 activity, respectively. (d, All population, *n* = 1,917 cells; with CDK2 activity, *n* = 1,038 cells; w/o CDK2 activity, *n* = 879 cells; CDK4/6^low^ cells, *n* = 4,127 cells. e, All population, *n* = 825 cells; with CDK2 activity, *n* = 545 cells; w/o CDK2 activity, *n* = 280 cells; CDK4/6^low^ cells, *n* = 2,333 cells. f, g, All population, *n* = 494 cells; with CDK2 activity, *n* = 332 cells; w/o CDK2 activity, *n* = 162 cells; CDK4/6^low^ cells, *n* = 1,253 cells). **(h)** Averaged cyclin E2 protein levels aligned to CDK4/6 activation in MCF-10A cells. Black, blue, and red lines correspond to all cells, cells with CDK2 activity, and cells without CDK2 activity, respectively. Dotted line indicates cyclin E2 protein level in CDK4/6^low^ cells. Data are mean ± 95% confidence interval (All population, *n* = 1,930 cells; with CDK2 activity, *n* = 1,226 cells; w/o CDK2 activity, *n* = 704 cells; CDK4/6^low^ cells, *n* = 3,986 cells). **(i)** Schematic of TMP-induced rapid expression of cyclin E1. **(j)** Averaged CDK2 activity traces aligned to the time of CDK4/6 activation for CDK4/6^high^ cells. Green dotted line indicates CDK4/6 activity onset. Data are mean ± 95% confidence interval (siRNA: Control, *n* = 3,172 cells; cyclin E1/2, *n* = 1,042 cells; DHFR-cyclin E1: DMSO, *n* = 470 cells; TMP, *n* = 932 cells). **(k)** Averaged E2F1 mRNA levels aligned to the time of CDK4/6 activation for CDK4/6^high^ cells. Dotted lines indicate E2F1 mRNA level in CDK4/6^low^ cells for each treatment. Data are mean ± 95% confidence interval (siRNA: Control, *n* = 1,699 cells; cyclin E1/2, *n* = 644 cells; DHFR-cyclin E1: DMSO, *n* = 233 cells; TMP, *n* = 552 cells). (j, k) Left, MCF-10A cells were transfected with control (black) or cyclin E1/2 siRNA (red) and were mitogen-released for various durations (9, 10, and 11 hr) prior to fixation. Right, MCF-10A cells expressing DHFR-cyclin E1 were mitogen-released for various durations (9, 10, and 11 hr) with DMSO (black) or TMP (50 µM, red) prior to fixation.

When we aligned live-cell traces and fixed data by the time of CDK4/6 activation (CDK4/6-on), we found heterogeneous responses in the timing of CDK2 activation (CDK2-on) (**Fig. 3b** and **Supplementary Fig. 3c**). Approximately 40% of CDK4/6^high^ cells did not activate CDK2 before fixation (**Supplementary Fig. 3d**). Thus, to test how CDK2 activation influences the kinetics of Rb phosphorylation, we further classified CDK4/6^high^ cells into populations with or without CDK2 activation (**Fig. 3c** and **Supplementary Fig. 3e–g**). When we plotted the percentage of Rb phosphorylation after CDK4/6 activation, we found that Rb phosphorylation at S807/811, T373, S608, and S780 were almost indistinguishable in CDK4/6^high^ cells regardless of CDK2 activity, indicating that CDK4/6 induces either multiple mono- or hyper-phosphorylation of Rb (**Fig. 3d–g**). However, when we measured E2F activity, CDK4/6^high^ cells with CDK2 activity had higher levels of E2F-target genes, E2F1, Cdc25A, and cyclin E2 mRNA and cyclin E2 protein (**Fig. 3h** and **Supplementary Fig. 4**). These data suggest two possibilities, that CDK2 activity facilitates E2F activity, or that heterogeneous E2F-activity levels correlate with the CDK2 classification, as E2F-target genes activate CDK2. To test these possibilities, we manipulated CDK2 activity and measured the kinetics of E2F activity. To modulate CDK2 activity, we used cyclin E1/2 siRNA to decrease endogenous cyclin E1/2 mRNA and the dihydrofolate reductase (DHFR)-trimethoprim (TMP) protein stabilization system^36^ to acutely increase exogenous cyclin E1 protein. In the absence of TMP, cyclin E1 protein conjugated with the DHFR domain is continuously degraded by the proteasome. Addition of TMP rapidly stabilizes cyclin E1 protein (**Fig. 3i**). We first found that knockdown of cyclin E1/2 and overexpression of cyclin E1 in CDK4/6^high^ cells delayed and accelerated CDK2 activation, respectively (**Fig. 3j** and **Supplementary Fig. 5a, b**). Notably, mRNA expression kinetics of E2F1 and Cdc25A after CDK4/6 activation were similar in CDK4/6^high^ cells regardless of CDK2 activity (**Fig. 3k** and **Supplementary Fig. 5c, d**). Taken together, the kinetics of Rb phosphorylation and mRNA levels of E2F target genes suggest that Rb phosphorylation by CDK4/6 activity is sufficient to initiate E2F induction.

**Figure 4.**
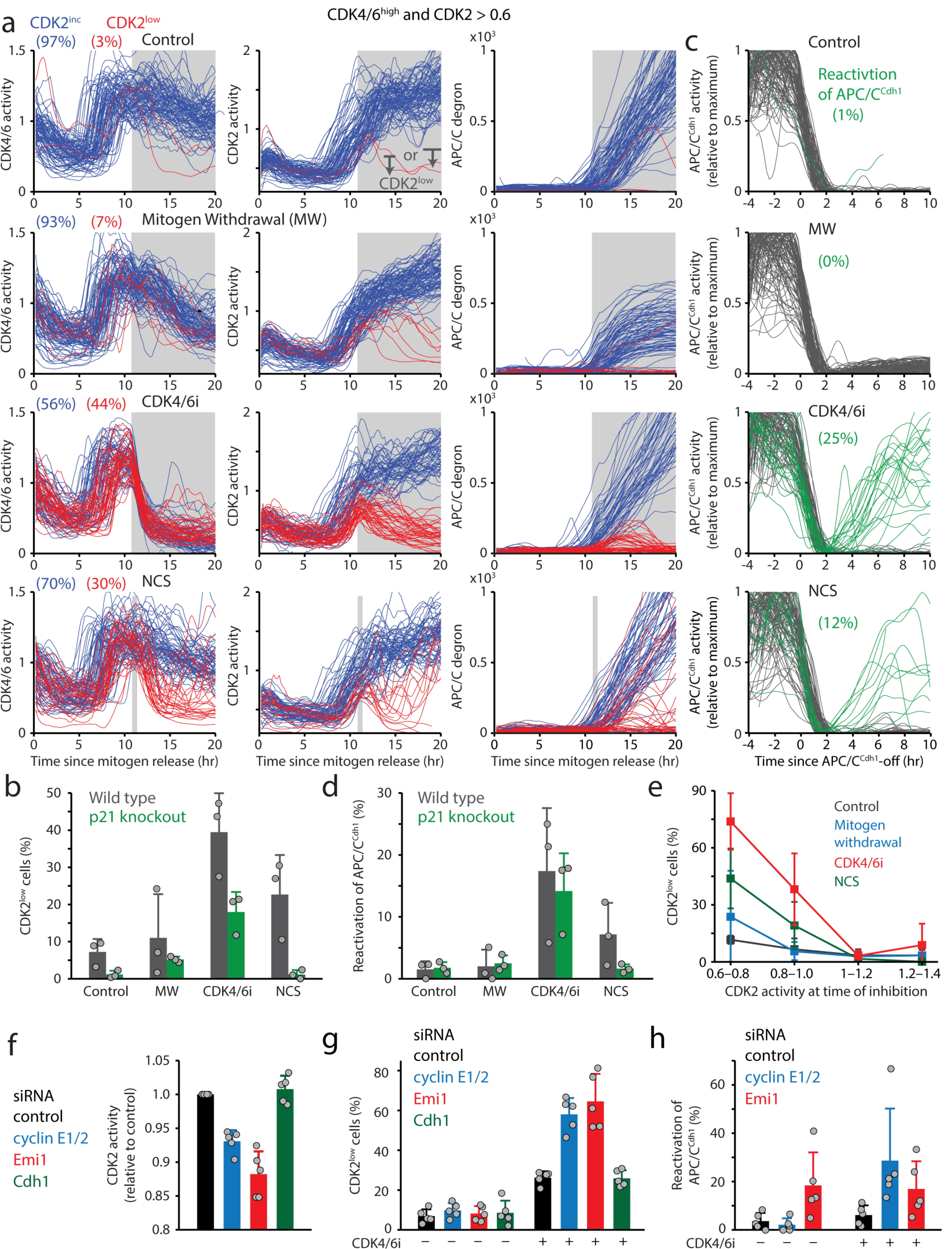
CDK2 activity level determines the commitment point in cell-cycle entry. **(a)** Single-cell traces of CDK4/6 activity (left), CDK2 activity (middle), and APC/C degron (right) in MCF-10A cells. Top to bottom, corresponding conditions under control, mitogen withdrawal, CDK4/6 inhibitor (palbociclib, 1 µM), and 10 min NCS pulse (100 ng/ml). All treatments performed at 11 hr (marked in gray). Blue and red lines correspond to CDK2^inc^ and CDK2^low^ cells, respectively. Cells were classified based on CDK2 activity after 4 and 9 hr of inhibition. **(b)** Percentages of CDK2^low^ cells in wild type (gray) and p21 knockout (green) MCF-10A cells for each treatment condition in (a). Data are mean ± s.d. (*n* = 3 biological replicates). **(c)** Single-cell traces of APC/C^Cdh1^ activity aligned to the time of APC/C^Cdh1^ inactivation in MCF-10A cells. Green traces correspond to cells reactivating APC/C^Cdh1^. **(d)** Percentage of cells reactivating APC/C^Cdh1^ in wild type (gray) and p21 knockout (green) MCF-10A cells for each treatment condition in (a). Data are mean ± s.d. (*n* = 3 biological replicates). **(e)** Percentages of CDK2^low^ cells for increasing levels of CDK2 activity at the time of inhibition, in wild type MCF-10A cells. Cells were classified based on each treatment condition in (a). Data are mean ± s.d. (*n* = 3 biological replicates). **(f)** Relative CDK2 activity for each knockdown condition as indicated in MCF-10A cells compared with control siRNA treated cells. Data are mean ± s.d. (*n* = 5 biological replicates). **(g, h)** Percentage of CDK2^low^ cells (g) and APC/C^Cdh1^ reactivating cells (h) for each knockdown condition as indicated in MCF-10A cells without and with CDK4/6 inhibitor (palbociclib, 1 µM). Data are mean ± s.d. (*n* = 5 biological replicates).

**Figure 5.**
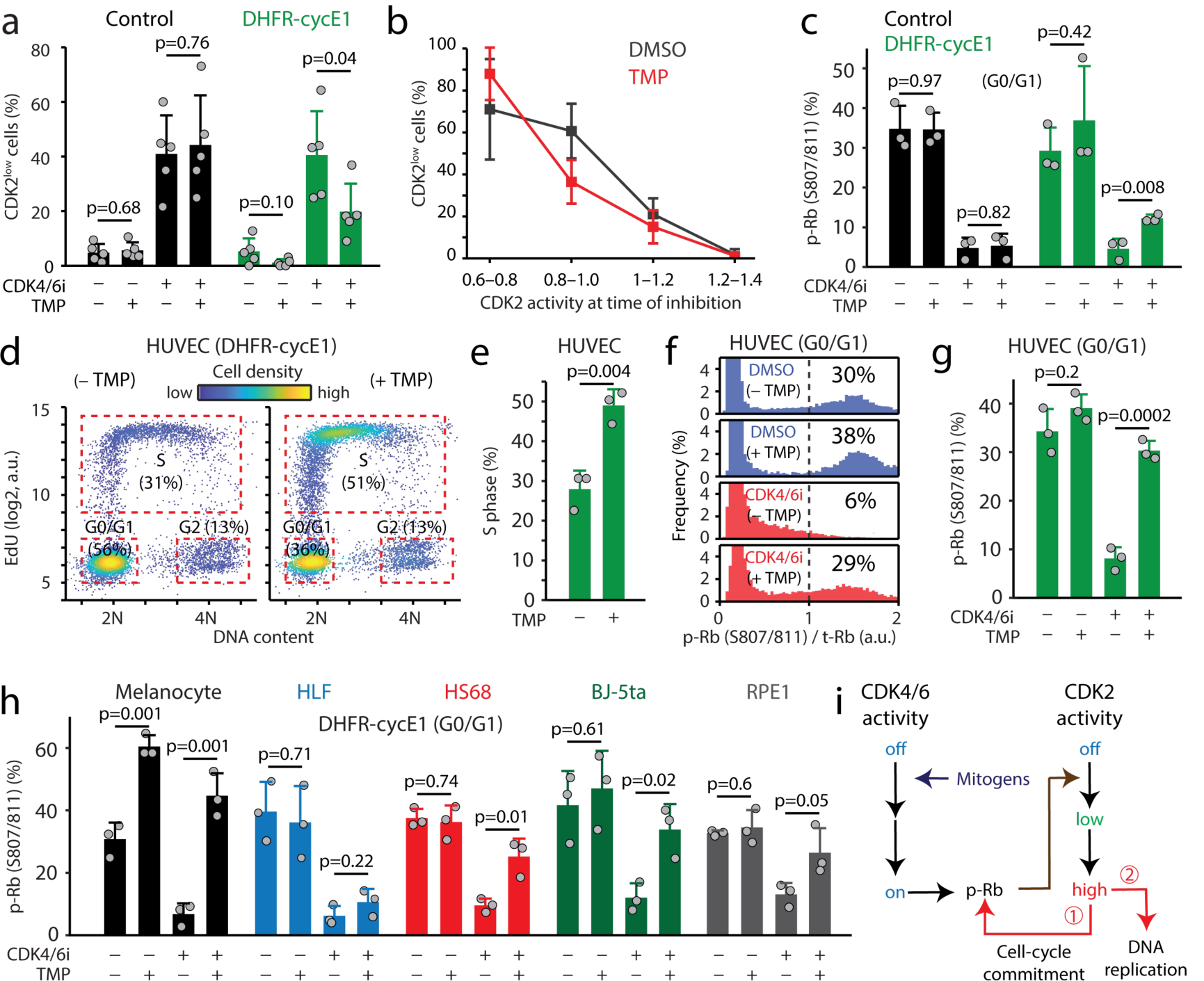
CDK2 activity sequentially regulates the commitment point and S-phase entry. **(a)** Percentage of CDK2^low^ cells for each condition as indicated in MCF-10A cells without and with CDK4/6 inhibitor (palbociclib, 1 µM). Data are mean ± s.d. (*n* = 5 biological replicates). **(b)** Percentages of CDK2^low^ cells for increasing levels of CDK2 activity at the time of inhibition in MCF-10A cells expressing DHFR-cycE1 without (black) or with TMP (50 µM, red) treatment for 6 hr. Data are mean ± s.d. (*n* = 5 biological replicates). **(c)** Percentage of p-Rb at S807/811 for each treatment condition as indicated for MCF-10A cells without (black) and with DHFR-cyclin E1 expression (green). **(d)** Scatterplot of Hoechst versus EdU, where cell density is color-coded. Cycling HUVECs expressing DHFR-cyclin E1 were treated with or without TMP (50 µM) for 6 hr followed by EdU (10 µM) treatment for 15 min prior to fixing and staining with Hoechst. **(e)** Percentage of cells in S phase for cycling HUVECs expressing DHFR-cyclin E1 with or without TMP (50 µM) treatment for 6 hr. Data are mean ± s.d. (*n* = 3 biological replicates). *P*-values were calculated with two-sample *t*-tests. **(f)** Histogram of p-Rb at S807/811 for HUVEC cells expressing DHFR-cyclin E1 in G0/G1 phase. Data are mean ± s.d. (*n* = 3 biological replicates). Cycling cells were treated with or without TMP (50 µM) for 6 hr followed by acute treatment with CDK4/6 inhibitor (palbociclib, 1 µM) and EdU (10 µM) for 15 min prior to fixation. **(g, h)** Percentage of p-Rb at S807/811 for each treatment condition as indicated in (g) for Melanocyte, HLF, HS68, BJ-5ta, and RPE1 cells (h) expressing DHFR-cyclin E1. Data are mean ± s.d. (*n* = 3 biological replicates). Cycling cells were treated with or without TMP (50 µM) for 6 hr followed by acute treatment with CDK4/6 inhibitor (palbociclib, 1 µM) and EdU (10 µM) for 15 min prior to fixation. *P*-values were calculated with two-sample *t*-tests. **(i)** Schematic of sequential regulation of the Rb/E2F pathway by CDK4/6 and CDK2. Rb phosphorylation by CDK4/6 induces E2F activity and gradual increase in CDK2 activity. High CDK2 activity triggers the commitment point by maintaining Rb phosphorylation, then initiates DNA replication.

### CDK2 activity determines cell-cycle progression in response to mitogen removal and DNA damage

We investigated when cell-cycle entry becomes CDK4/6 independent and how mitogenic and DNA damage signaling regulate CDK4/6 and CDK2 activities regarding the commitment point. To test this, we withdrew mitogens and then applied the CDK4/6 inhibitor (palbociclib) or induced DNA damage signaling 11 hours after mitogen release. For analysis, we selected CDK4/6^high^ cells with CDK2 activity (>0.6) at the time of inhibition. Based on CDK2 response to the treatment, we used two windows to further classify cells by path, either continuing to increase CDK2 activity for proliferation (CDK2^inc^) or reversing CDK2 activation and entering quiescence (CDK2^low^) (**Fig. 4a**). Mitogen withdrawal inhibited CDK4/6 activity with a 4–5 hour delay and only slightly increased the percentage of CDK2^low^ cells (**Fig. 4a, b**). To induce DNA damage signaling, we incubated cells with neocarzinostatin (NCS) for 10 minutes to produce exogenous DNA double-stranded breaks. Both palbociclib and NCS rapidly suppressed CDK4/6 activity and significantly increased the percentage of CDK2^low^ cells (**Fig. 4a, b**). We note that CDK4/6 activity was not inhibited by NCS in CDK2^inc^ cells. As CDK2^inc^ cells may have already entered S phase at the time of drug treatment, it is plausible that this reduced sensitivity of CDK4/6 activity to NCS in CDK2^inc^ cells is due to the absence of p21, a CDK inhibitor that is degraded during S phase^37–39^. Consistently, our findings showed that NCS-mediated inactivation of CDK4/6 and consequent reversibility in cell-cycle entry were absent in p21 knockout cells (**Fig. 4b** and **Supplementary Fig. 6a**).

In these experiments, we noticed that APC/C^Cdh1^ inactivation was reversed by CDK4/6 inhibition in both wild type and p21 knockout cells (**Fig. 4a** and **Supplementary Fig. 6b, c**). When we examined the accumulation rate of the reporter for APC/C^Cdh1^ activity^8^, CDK4/6 inhibition caused reactivation of APC/C^Cdh1^ and cell-cycle exit (**Fig. 4c, d** and **Supplementary Fig. 6b, c**). Recent work using fission yeast showed that different CDK substrates get phosphorylated at different levels of CDK activity^40^. Additional studies in mammalian cells demonstrated that high CDK2 activity is required to initiate CDK2-Rb feedback^24^. Given these studies, we next tested whether a threshold level of CDK2, instead of APC/C^Cdh1^ inactivation, correlates with irreversible cell-cycle entry. We consistently found that the level of CDK4/6 activity typically plateaued before CDK4/6 inhibition and was not correlated with following CDK2 activity (**Supplementary Fig. 6d, e**). In addition, our data show that CDK2 activity predicts cell-cycle entry or exit in both wild type and p21 knockout cells at the time of CDK4/6 inhibition (**Fig. 4e** and **Supplementary Fig. 6f**). When CDK2 activity was over 1, the level at which DNA replication has previously been reported to begin^10^, cells likely continued through cell-cycle entry regardless of CDK4/6 activity and changes in external signaling. These findings suggest that CDK2 activity level is the primary determinant to trigger irreversible cell-cycle commitment with respect to mitogenic and DNA damage signaling.

We next hypothesize that APC/C^Cdh1^ inactivation controls CDK2 activity by regulating cyclin A protein degradation^41,42^ and may consequently potentiate cell-cycle commitment. Thus, we further tested how APC/C^Cdh1^ inhibition regulates the commitment point. To modulate APC/C^Cdh1^ activity, we used Cdh1 (APC/C coactivator) and Emi1 (APC/C inhibitor) siRNA to prematurely inactivate APC/C^Cdh1^ and to delay APC/C^Cdh1^ inactivation, respectively. As a control, we used cyclin E1/2 siRNA to reduce CDK2 activity. To test the percentage of cell that cross the commitment point with respect to CDK4/6 inhibition, we selected CDK4/6^high^ cells with CDK2 activity (>0.6) 11 hours after mitogen release. When we compared CDK2 activity at the time of CDK4/6 inhibition, we found the Emi1 and cyclin E1/2 knockdown condition showed lower levels of CDK2 activity compared to the control condition (**Fig. 4f**). Cdh1 knockdown showed higher basal intensity of the APC/C reporter compared to the control as APC/C^Cdh1^ activity was decreased (**Supplementary Fig. 6g**). Accordingly, after CDK4/6 inhibition, our data show that Emi1 knockdown increased the percentage of CDK2^low^ cells to almost the same level as cyclin E1/2 knockdown (**Fig. 4g**). However, since inactivation of APC/C^Cdh1^ by Cdh1 knockdown did not further increase CDK2 activity (**Fig. 4f**), the percentage of CDK2^low^ cells in the Cdh1-knockdown condition was similar to that of the control (**Fig. 4g**). Intriguingly, while Emi1 knockdown increased the percentage of cells reactivating APC/C^Cdh1^ regardless of CDK4/6 inhibition, cyclin E1/2 knockdown increased the percentage of cells reactivating APC/C^Cdh1^ after CDK4/6 inhibition (**Fig. 4h**). Our data suggest that APC/C^Cdh1^ inactivation contributes to cell-cycle commitment likely through CDK2 activity.

### CDK2 activity coordinates the timing of CDK4/6-independent Rb phosphorylation and sequential DNA replication

Previous studies showed that CDK2 activation by cyclin E overexpression can bypass CDK4/6 activity^43–47^. Using the DHFR-cyclin E1 protein stabilization system^36^, we increased CDK2 activity and further tested whether CDK2 activity at the time of CDK4/6 inhibition is able to predict cell-cycle commitment. We added palbociclib 11 hours after mitogen release with or without TMP, and selected CDK4/6^high^ cells with CDK2 activity (>0.6). We found that addition of TMP did not alter either CDK2 activity or the percentage of CDK2^inc^ cells in the control condition, in which cells did not express the DHFR-cyclin E1 construct (**Fig. 5a** and **Supplementary Fig. 7a**). While we confirmed that increased CDK2 activity led to a higher percentage of CDK2^inc^ cells, we found that CDK2 activity at the time of CDK4/6 inhibition could predict cell-cycle entry or exit in cells with or without cyclin E1 overexpression (**Fig. 5a, b** and **Supplementary Fig. 7b**). Furthermore, after CDK4/6 inhibition, the percentage of cells reactivating APC/C^Cdh1^ was significantly lower when we increased CDK2 activity (**Supplementary Fig. 7c**). Our data indicate that a threshold level of CDK2 activity is the key regulator controlling cell-cycle commitment.

We next sought to determine the molecular mechanism of CDK2-mediated Rb phosphorylation. Previous reports demonstrated that CDK2 activity controls not only Rb phosphorylation, but also the initiation and progression of DNA replication^48–50^. A recent study also showed that high CDK2 can initiate Rb phosphorylation at the onset of S phase^24^. This can be explained by two possibilities, that the timing of Rb phosphorylation and DNA replication by CDK2 activity is tightly coupled, or that other mechanisms in S phase are required for CDK2 to induce Rb phosphorylation. To address this, we tested whether an increase in CDK2 activity is enough to induce Rb phosphorylation before DNA replication. To exclude the possibility of contribution by other regulatory mechanisms during S phase, we used 5-ethynyl-20-deoxyuridine (EdU) and Hoechst staining to gate cells in G0/G1. During G0/G1, while control cells almost completely lost Rb phosphorylation at S807/811 following acute CDK4/6 inhibition for 15 min, an increase in CDK2 activity by cyclin E1 overexpression significantly upregulated the percentage of Rb phosphorylation at S807/811 in MCF-10A cells (**Fig. 5c** and **Supplementary Fig. 8a**). Furthermore, an increase in CDK2 activity accelerated S-phase entry (**Supplementary Fig. 8b**), suggesting that high CDK2 activity coordinates the timing of CDK2-Rb feedback and DNA replication. We next tested the effect of cyclin E1 overexpression on initiation of CDK2-Rb feedback and DNA replication in asynchronously cycling cells. When we increased CDK2 activity by addition of TMP in passage-limited human umbilical vein endothelial cells (HUVECs) expressing DHFR-cyclin E1 for 6 hours, the percentage of cells entering S phase was significantly increased (**Fig. 5d, e**), confirming that CDK2 activity is the limiting factor to induce S-phase entry in asynchronously cycling cells. In addition, an increase in CDK2 activity significantly rescued the reduction of Rb phosphorylation at S807/811 in G0/G1 cells by acute CDK4/6 inhibition (**Fig. 5f, g**), which suggests that high CDK2 activity is sufficient to maintain Rb phosphorylation. We observed similar results that CDK2 sequentially regulates the timing of CDK2-Rb feedback and S-phase entry in passage-limited human melanocytes, and HS68, as well as hTERT immortalized BJ-5ta and retinal pigment epithelial (RPE1-hTERT) cells, but not in HLF (**Fig. 5h** and **Supplementary Fig. 8c**). These data indicate that CDK2-mediated regulation of irreversible cell-cycle entry and S-phase entry is conserved across primary and immortalized cell types. In the case of CDK2 knockout cells^51,52^, it has been reported that CDK1 binds to cyclin E and compensates for the loss of CDK2 by phosphorylating CDK2 substrates^8,53^. Taken together, our data propose that high CDK2 activity coordinates the timing of irreversible cell-cycle entry and transition into S phase.

## Discussion

Our studies show that CDK4/6 activity initiates E2F activation and high CDK2 activity coordinates the timing of the commitment point and DNA replication in mitogen-starved and cycling cells (**Fig. 5i**). Furthermore, we propose that CDK2-Rb feedback is the primary signaling network regulating cell-cycle commitment. Given APC/C^Cdh1^ degrades cyclin A protein, APC/C^Cdh1^ inactivation likely further increases CDK2 activity and potentiates engagement of the CDK2-Rb pathway. Other regulatory components downstream of Rb phosphorylation, including positive feedback from Skp2 autoinduction^54–57^ and E2F autoregulation^58,59^, have been also proposed as regulators of the commitment point. Although our study did not directly investigate these pathways, it is possible that they also contribute to cell-cycle commitment by controlling CDK2 activity (**Supplementary Fig. 8d**). While our approach is limited in differentiating between mono-phosphorylation and hyper-phosphorylation of Rb, our data indicate that Rb phosphorylation by CDK4/6 activity is tightly correlated with the initiation of E2F transcription activity before CDK2 activation, suggesting that Rb phosphorylation by CDK4/6 activity is sufficient to trigger E2F activation. Therefore, it is likely that CDK4/6 could mono-phosphorylate Rb and induce E2F activity^19,23^ or trigger mono- and hyper-phosphorylation of Rb and E2F activity^19,24^.

Our studies examined the short-term effect of CDK4/6 inhibitors (15 min – 5 hours). A recent literature demonstrated that long-term treatment with CDK4/6 inhibitors (48 hours) can cause cell-cycle arrest by redistributing CDK inhibitor proteins, notably p21^(35)^. Since high CDK2 activity is required to phosphorylate Rb, the loss of Rb phosphorylation by the short-term treatment with CDK4/6 inhibitors is likely on-target inhibition of E2F-mediated cyclin E expression. In addition, although CDK4/6 inhibitors induced redistribution of other CDK inhibitor proteins, p27 and p57, which can inhibit CDK2 activity, we showed that CDK4/6 inhibitors suppressed low CDK2 activity and induced cell-cycle arrest in p21 knockout MCF-10A cells.

Consistent with a recent study^24^, our studies demonstrate that high CDK2 activity is required for CDK2 to phosphorylate Rb in mitogen-starved an cycling cells. Therefore, a threshold level of CDK2 activity regulates the commitment point by initiating Rb phosphorylation. A similar concept, proposed by the Jan Skotheim laboratory, suggests that a CDK activity threshold in cycling cells determines passage through the commitment point^32^. Furthermore, our study and recent other studies propose that mitogen removal and various stress induction cause rapid and slow inactivation of CDK4/6, respectively^9,24^. Depending on the strength of anti-mitogens, different external stimuli may reverse CDK4/6 activity and Rb phosphorylation with distinct kinetics. Consequently, individual external stimuli could reverse cell-cycle entry at different time points before a threshold level of CDK2 activity which induces Rb phosphorylation and DNA replication. This provides insights into the control system and safety mechanism for cells to 1) withdraw from cell-cycle entry depending on external conditions, and 2) protect against incomplete DNA replication.

## METHODS

Methods and associated references are available in the online version of the paper.

## ACKNOWLEDGMENTS

We thank Nalin Ratnayeke for a pre-extraction protocol; Mingyu Chung for helpful comments; Tobias Meyer for helpful discussion and support; Richard Baer, Michele Pagano, and Charles Sherr for helpful discussions and critical comments. This work was supported by the Neuroendocrine Tumor Research Foundation (M.K., 653132) and Herbert Irving Comprehensive Cancer Center (H.Y., P30 CA013696).

## AUTHOR CONTRIBUTIONS

S.K., A.L, C.N., H.Y., and M.K. designed the research and performed experiments. S.K., A.L, H.Y., and M.K. wrote the manuscript discussed with all authors.

## METHODS

### Cell Culture

MCF-10A human mammary epithelial cells (ATCC, CRL-10317) were cultured in phenol red-free DMEM/F12 (Invitrogen) and supplemented with 5% horse serum, 20 µg/ml EGF, 10 µg/ml insulin, 500 µg/ml hydrocortisone, 100 ng/ml cholera toxin, 50 U/ml penicillin, and 50 µg/ml streptomycin. For mitogen starvation, DMEM/F12 plus 0.3% BSA w/v, 500 µg/ml hydrocortisone, and 100 ng/ml cholera toxin was used. HLF primary human lung fibroblasts (Cell Appliance, 506K-05f) were cultured in HLF Growth Medium (Cell Appliance, 516K-500) kit. Human primary epidermal melanocytes (ATCC, PCS-200-013) were culture in Dermal Cell Basal Medium (PCS-200-030). HS68 primary human foreskin fibroblasts (ATCC, CRL-1635) were cultured in DMEM (Invitrogen) plus 10% FBS, 50 U/ml penicillin, and 50 µg/ml streptomycin. HUVECs human umbilical vein endothelial cells (Lonza, C2519A) were cultured in Endothelial Cell Growth Medium (Promo Cell, C-22010) kit. BJ-5ta human foreskin fibroblasts (ATCC, CRL-4001) were cultured in DMEM (Invitrogen) plus 10% FBS, 50 U/ml penicillin, and 50 µg/ml streptomycin. RPE1-hTERT human retinal pigment epithelial cells (ATCC, CRL-4000) were cultured in DMEM: F12 (Invitrogen) plus 10% FBS and 0.01 mg/mL hygromycin B. All cell lines tested negative for mycoplasma.

### Antibodies and Reagents

Palbociclib (S1116), Abemaciclib (S7158), Ribociclib (S7440), and Roscovitine (S1153) were purchased from Selleck Chemicals. Neocarzinostatin (N9162) was from Sigma-Aldrich. RO-3306 (217721) and CDK2i III (CVT-315, 238803) were from Millipore. Anti-phosphor-Rb (Ser807/811) (8516) and anti-phosphor-Rb (S608) (2181) were purchased from Cell Signaling Technology. Anti-phosphor-Rb (S780) (558385) and anti-Rb (554136) were purchased from BD Bioscience, and anti-phophor-Rb (T373) (ab52975) was obtained from Abcam.

### Constructs and stable cell lines

pLenti-DHB(a.a.994–1087)-mVenus-p2a-mCherry-Rb (a.a.886–928) and mCerulean-Geminin (a.a.1–110)-p2a-H2B-iRFP670 were described previously^9^. Full sequence and constructs are available from Addgene. DHFR-cyclin E1 was cloned into a pCru5 retroviral vector. To generate stable cell lines, lentiviral and retroviral constructs were introduced into MCF-10A, HLF, melanocyte, HS68, BJ-5ta, and RPE1-hTERT cells by viral transduction.

### Immunofluorescence

Cells were fixed by adding 4% paraformaldehyde at a ratio of 1:1 to culture medium (final 2% paraformaldehyde) for 15 min. Then, cells were washed three times in PBS, followed by incubation in permeabilization/blocking with 0.1% triton X-100, 10% FBS, 1% BSA, and 0.01% NaN3 for 1 hr, and stained overnight at 4°C with primary antibodies. Primary antibodies were visualized using a secondary antibody conjugated to Alexa Fluor-488, -568, or -647. For EdU staining, cells were treated with 10 µM EdU for 15 min, then fixed and processed with aziede-modified Alexa Fluor-647 according to manufacturer’s instructions (Invitrogen, #C10269). To prevent the use of fluorophores limited by fluorescent reporters, cells were chemically bleached^60^. Pre-extraction^61^ and multiplexed imaging^34^ were previously described.

### RNA FISH

RNA *in situ* hybridization was carried out using the Affymetrix Quantigene ViewRNA ISH cell assay as specified in the user manual. Briefly, cells were plated in a 96-well glass plate (Cellvis P96-1.5H-N) that was pre-hybridized with collagen (Advanced BioMatrix, #5005-B) 1:100 in PBS overnight. At the time of fixing, cells were fixed with 4% paraformaldehyde for 15 min and dehydrated overnight using 75% EtOH. After rehydration in PBS for 10 min, the cells were permeabilized with 0.2% TX-100 for 15min RT, and then treated for probe hybridization, amplification, and labeling with Alexa Fluor 555. Cells were then incubated with Hoechst (1:10,000 in PBS) for 10 min, washed three times with PBS, and left in PBS for imaging. For the cases where immunofluorescence was to be additionally performed, after imaging the FISH signal, cells were incubated with the ViewRNA wash buffer as specified in the user manual to remove the probes and allow for measurement of other fluorophores (Alexa 488 and Alexa 647).

### siRNA Transfection

siRNA was transfected using DharmaFECT1 Transfection Reagent (Horizon, T-2001) according to the manufacturer’s instructions. The following siRNAs were used: Control (Integrated DNA Technologies (IDT)), Negative Control siRNA, CCNE1 (Dharmacon, siGENOME pool), CCNE2 (Dharmacon, siGENOME pool), CDH1 (IDT, hs.Ri.FZR1.13.1-3), Emi1 (IDT, hs.Ri.FBXO5.13.1-3).

### CDK4 correction based on CDK2 reporter

Based on data from CDK inhibitors and MEF lacking cyclin E1/2 and A1/2, we previously reported the contribution of CDK2 activity to the CDK4/6 reporter^9^. To correct for the CDK2 contribution, we applied the correction factor described previously^9^. Briefly, in the presence of CDK4/6 inhibitor, we used linear regression to calculate the contribution of CDK2 activity to the CDK4/6 reporter. CDK4/6 activity = CDK4/6 activity – 35% × CDK2 activity

### Microscopy

All images were taken on an Axio Observer 7 microscope (Zeiss) using 20X objective (0.8 N.A) with 2-by-2 pixel binning. For live-cell imaging, cells were imaged every 12 min in a humidified and 37°C chamber in 5% CO_2_. Total light exposure time was kept under 400 ms for each time point.

### Image Analysis

All image analyses were performed with custom Matlab scripts as previously described^8,62^. Briefly, after illumination bias correction, cells were segmented, either by using Hoechst staining (fixed-cell imaging) or H2B-iRPF670 (live-cell imaging). For DHB-mVenus and mCherry-Rb measurements, cells were segmented for their cytoplasmic regions by spatially approximating a ring with an inner radius of 2 µm outside of the nuclear mask and an outer radius maximum of 10 µm outside the nuclear mask. Regions within 10 µm of another nucleus were excluded. Regions with pixel intensities indistinguishable from background (discussed below) were also excluded. For RNA FISH measurements, cells were segmented for their whole cell regions by using an area that encompasses the nucleus and reaches out as far as 50 µm outside of the nuclear mask while preventing overlap with neighboring cells. A mask of FISH puncta was generated by top hat-filtering raw images with a circular kernel of radius 4 µm and thresholding absolute intensity. The RNA puncta parameter represents an average of the number of pixels in whole cell regions. For cell tracking, the deflection-bridging algorithm was implemented to perform tracking of cells between live-cell frames as well as between the final live-cell frame and subsequent fixed-cell image.

### Statistical Analysis

Statistical analyses were based on a two-sample *t*-test. The data were represented either mean ± s.d. or mean ± 95% confidence intervals and the number of replicates was indicated in respective figure legends. *P*-values were indicated in respective figures. No statistical methods were used to predetermine sample size. The experiments were not randomized and the investigators were not blinded to allocation during experiments and outcome assessment.

## Data Availability

The data that support the findings of this study are available within the paper and Supplementary Information files. Extra data are available from the corresponding authors upon request.

## Code availability

The code for the image analysis pipeline is available at https://github.com/scappell/Cell_tracking. Additional modified scripts and source data are available from the corresponding authors upon request.

## REFERENCES

1. Hanahan, D. & Weinberg, R. A. The hallmarks of cancer. Cell 100, 57–70 (2000).

2. Vincent, I., Pae, C. I. & Hallows, J. L. The cell cycle and human neurodegenerative disease. Prog. Cell Cycle Res. 5, 31–41 (2003).

3. Henley, S. A. & Dick, F. A. The retinoblastoma family of proteins and their regulatory functions in the mammalian cell division cycle. Cell Div. 7, 10 (2012).

4. Matson, J. P. & Cook, J. G. Cell cycle proliferation decisions: the impact of single cell analyses. FEBS J. 284, 362–375 (2017).

5. Pardee, A. B. A restriction point for control of normal animal cell proliferation. Proc. Natl. Acad. Sci. U. S. A. 71, 1286–90 (1974).

6. Erol, A. Genotoxic stress-mediated cell cycle activities for the decision of cellular fate. Cell Cycle 10, (2011).

7. Deckbar, D., Jeggo, P. A. & Löbrich, M. Understanding the limitations of radiation-induced cell cycle checkpoints. Crit. Rev. Biochem. Mol. Biol. 46, 271–83 (2011).

8. Cappell, S. D., Chung, M., Jaimovich, A., Spencer, S. L. & Meyer, T. Irreversible APCCdh1 Inactivation Underlies the Point of No Return for Cell-Cycle Entry. Cell 166, 167–180 (2016).

9. Yang, H. W. et al. Stress-mediated exit to quiescence restricted by increasing persistence in CDK4/6 activation. Elife 9, (2020).

10. Spencer, S. L. et al. The proliferation-quiescence decision is controlled by a bifurcation in CDK2 activity at mitotic exit. Cell 155, 369–383 (2013).

11. Cappell, S. D. et al. EMI1 switches from being a substrate to an inhibitor of APC/CCDH1 to start the cell cycle. Nature 558, 313–317 (2018).

12. Lee, J. O., Russo, A. A. & Pavletich, N. P. Structure of the retinoblastoma tumour-suppressor pocket domain bound to a peptide from HPV E7. Nature 391, 859–865 (1998).

13. Fisher, R. P. Getting to S: CDK functions and targets on the path to cell-cycle commitment. F1000Research 5, 2374 (2016).

14. Weinberg, R. A. The retinoblastoma protein and cell cycle control. Cell 81, 323–30 (1995).

15. Lundberg, A. S. & Weinberg, R. A. Functional inactivation of the retinoblastoma protein requires sequential modification by at least two distinct cyclin-cdk complexes. Mol. Cell. Biol. 18, 753–61 (1998).

16. Connell-Crowley, L., Harper, J. W. & Goodrich, D. W. Cyclin D1/Cdk4 regulates retinoblastoma protein-mediated cell cycle arrest by site-specific phosphorylation. Mol. Biol. Cell 8, 287–301 (1997).

17. Matsushime, H. et al. Identification and properties of an atypical catalytic subunit (p34PSK-J3/cdk4) for mammalian D type G1 cyclins. Cell 71, 323–334 (1992).

18. Kato, J., Matsushime, H., Hiebert, S. W., Ewen, M. E. & Sherr, C. J. Direct binding of cyclin D to the retinoblastoma gene product (pRb) and pRb phosphorylation by the cyclin D-dependent kinase CDK4. Genes Dev. 7, 331–342 (1993).

19. Narasimha, A. M. et al. Cyclin D activates the Rb tumor suppressor by mono-phosphorylation. Elife 2014, 1–21 (2014).

20. Burke, J. R., Hura, G. L. & Rubin, S. M. Structures of inactive retinoblastoma protein reveal multiple mechanisms for cell cycle control. Genes Dev. 26, (2012).

21. Burke, J. R., Deshong, A. J., Pelton, J. G. & Rubin, S. M. Phosphorylation-induced conformational changes in the retinoblastoma protein inhibit E2F transactivation domain binding. J. Biol. Chem. 285, 16286–16293 (2010).

22. Rubin, S. M. Deciphering the Rb phosphorylation code. Trends Biochem. Sci. 38, (2014).

23. Sanidas, I. et al. A Code of Mono-phosphorylation Modulates the Function of RB. Mol. Cell 73, 985–1000.e6 (2019).

24. Chung, M. et al. Transient Hysteresis in CDK4/6 Activity Underlies Passage of the Restriction Point in G1. Mol. Cell 76, 562–573 (2019).

25. Merrick, K. a. et al. Switching Cdk2 On or Off with Small Molecules to Reveal Requirements in Human Cell Proliferation. Mol. Cell 42, 624–636 (2011).

26. Castro, A., Bernis, C., Vigneron, S., Labbé, J. C. & Lorca, T. The anaphase-promoting complex: A key factor in the regulation of cell cycle. Oncogene 24, 314–325 (2005).

27. Sivakumar, S. & Gorbsky, G. J. Spatiotemporal regulation of the anaphase-promoting complex in mitosis. Nat. Publ. Gr. 16, 82–94 (2015).

28. Hsu, J. Y., Reimann, J. D. R., Sørensen, C. S., Lukas, J. & Jackson, P. K. E2F-dependent accumulation of hEmi1 regulates S phase entry by inhibiting APC(Cdh1). Nat. Cell Biol. 4, 358–66 (2002).

29. Miller, J. J. et al. Emi1 stably binds and inhibits the anaphase-promoting complex/cyclosome as a pseudosubstrate inhibitor. Genes Dev. 20, 2410–2420 (2006).

30. Hahn, A. T., Jones, J. T. & Meyer, T. Quantitative analysis of cell cycle phase durations and PC12 differentiation using fluorescent biosensors. Cell Cycle 8, 1044–52 (2009).

31. Sakaue-Sawano, A. et al. Visualizing Spatiotemporal Dynamics of Multicellular Cell-Cycle Progression. Cell 132, 487–498 (2008).

32. Schwarz, C. et al. A Precise Cdk Activity Threshold Determines Passage through the Restriction Point. Mol. Cell 69, 253–264.e5 (2018).

33. Min, M. & Spencer, S. L. Spontaneously slow-cycling subpopulations of human cells originate from activation of stress-response pathways. PLoS Biol. 17, (2019).

34. Gut, G., Herrmann, M. D. & Pelkmans, L. Multiplexed protein maps link subcellular organization to cellular states. Science 361, eaar7042 (2018).

35. Guiley, K. Z. et al. P27 allosterically activates cyclin-dependent kinase 4 and antagonizes palbociclib inhibition. Science (80-.). 366, eaaw2106 (2019).

36. Iwamoto, M., Björklund, T., Lundberg, C., Kirik, D. & Wandless, T. J. A General Chemical Method to Regulate Protein Stability in the Mammalian Central Nervous System. Chem. Biol. 17, 981–988 (2010).

37. Bunz, F. Requirement for p53 and p21 to Sustain G2 Arrest After DNA Damage. Science (80-.). 282, 1497–1501 (1998).

38. Krenning, L., Feringa, F. M., Shaltiel, I. A., van den Berg, J. & Medema, R. H. Transient Activation of p53 in G2 Phase Is Sufficient to Induce Senescence. Mol. Cell 55, 59–72 (2014).

39. Abbas, T. et al. PCNA-dependent regulation of p21 ubiquitylation and degradation via the CRL4Cdt2 ubiquitin ligase complex. Genes Dev. 22, (2008).

40. Swaffer, M. P., Jones, A. W., Flynn, H. R., Snijders, A. P. & Nurse, P. CDK Substrate Phosphorylation and Ordering the Cell Cycle. Cell 167, 1750–1761.e16 (2016).

41. Zachariae, W. & Nasmyth, K. Whose end is destruction: Cell division and the anaphase-promoting complex. Genes and Development 13, 2039–2058 (1999).

42. Sudakin, V. et al. The cyclosome, a large complex containing cyclin-selective ubiquitin ligase activity, targets cyclins for destruction at the end of mitosis. Mol. Biol. Cell 6, 185–198 (1995).

43. Lukas, J. et al. Cyclin E-induced S phase without activation of the pRb/E2F pathway. Genes Dev. 11, 1479–92 (1997).

44. Keenan, S. M., Lents, N. H. & Baldassare, J. J. Expression of Cyclin E Renders Cyclin D-CDK4 Dispensable for Inactivation of the Retinoblastoma Tumor Suppressor Protein, Activation of E2F, and G 1-S Phase Progression. J. Biol. Chem. 279, (2004).

45. Hinds, P. W. et al. Regulation of retinoblastoma protein functions by ectopic expression of human cyclins. Cell 70, 993–1006 (1992).

46. Duronio, R. J. & O’Farrell, P. H. Developmental control of the G1 to S transition in Drosophila: Cyclin E is a limiting downstream target of E2F. Genes Dev. 9, (1995).

47. Álvarez-Fernández, M. & Malumbres, M. Mechanisms of Sensitivity and Resistance to CDK4/6 Inhibition. Cancer Cell 37, (2020).

48. Fisher, D. Control of DNA replication by cyclin-dependent kinases in development. Results Probl. Cell Differ. 53, 201–17 (2011).

49. Tanaka, S., Tak, Y.-S. & Araki, H. The role of CDK in the initiation step of DNA replication in eukaryotes. Cell Div. 2, 16 (2007).

50. Krude, T., Jackman, M., Pines, J. & Laskey, R. A. Cyclin/Cdk-dependent initiation of DNA replication in a human cell-free system. Cell 88, 109–19 (1997).

51. Ortega, S. et al. Cyclin-dependent kinase 2 is essential for meiosis but not for mitotic cell division in mice. Nat. Genet. 35, 25–31 (2003).

52. Berthet, C., Aleem, E., Coppola, V., Tessarollo, L. & Kaldis, P. Cdk2 Knockout Mice Are Viable. Curr. Biol. 13, 1775–1785 (2003).

53. Aleem, E., Kiyokawa, H. & Kaldis, P. Cdc2-cyclin E complexes regulate the G1/S phase transition. Nat. Cell Biol. 7, (2005).

54. Amador, V., Ge, S., Santamaría, P. G., Guardavaccaro, D. & Pagano, M. APC/CCdc20 Controls the Ubiquitin-Mediated Degradation of p21 in Prometaphase. Mol. Cell 27, (2007).

55. Kossatz, U. et al. Skp2-dependent degradation of p27kip1 is essential for cell cycle progression. Genes Dev. 18, 2602–7 (2004).

56. Carrano, A. C., Eytan, E., Hershko, A. & Pagano, M. SKP2 is required for ubiquitin-mediated degradation of the CDK inhibitor p27. Nat. Cell Biol. 1, 193–9 (1999).

57. Yung, Y., Walker, J. L., Roberts, J. M. & Assoian, R. K. A Skp2 autoinduction loop and restriction point control. J. Cell Biol. 178, 741–7 (2007).

58. Yao, G., Lee, T. J., Mori, S., Nevins, J. R. & You, L. A bistable Rb-E2F switch underlies the restriction point. Nat. Cell Biol. 10, (2008).

59. Johnson, D. G., Ohtani, K. & Nevins, J. R. Autoregulatory control of E2F1 expression in response to positive and negative regulators of cell cycle progression. Genes Dev. 8, 1514–1525 (1994).

60. Lin, J.-R., Fallahi-Sichani, M. & Sorger, P. K. Highly multiplexed imaging of single cells using a high-throughput cyclic immunofluorescence method. Nat. Commun. 6, 8390 (2015).

61. HÅland, T. W., Boye, E., Stokke, T., Grallert, B. & SyljuÅsen, R. G. Simultaneous measurement of passage through the restriction point and MCM loading in single cells. Nucleic Acids Res. 43, e150 (2015).

62. Yang, H. W., Chung, M., Kudo, T. & Meyer, T. Competing memories of mitogen and p53 signalling control cell-cycle entry. Nature 549, 404–408 (2017).

